# The mechanisms behind perivascular fluid flow

**DOI:** 10.1101/2020.06.17.157917

**Authors:** Cécile Daversin-Catty, Vegard Vinje, Kent-André Mardal, Marie E. Rognes

## Abstract

Flow of cerebrospinal fluid (CSF) in perivascular spaces (PVS) is one of the key concepts involved in theories concerning clearance from the brain. Experimental studies have demonstrated both net and oscillatory movement of microspheres in PVS (Mestre et al. (2018), Bedussi et al. (2018)). The oscillatory particle movement has a clear cardiac component, while the mechanisms involved in net movement remain disputed. Using computational fluid dynamics, we computed the CSF velocity and pressure in a PVS surrounding a cerebral artery subject to different forces, representing arterial wall expansion, systemic CSF pressure changes and rigid motions of the artery. The arterial wall expansion generated velocity amplitudes of 60–260 *µ*m/s, which is in the upper range of previously observed values. In the absence of a static pressure gradient, predicted net flow velocities were small (<0.5 *µ*m/s), though reaching up to 7 *µ*m/s for non-physiological PVS lengths. In realistic geometries, a static systemic pressure increase of physiologically plausible magnitude was sufficient to induce net flow velocities of 20–30 *µ*m/s. Moreover, rigid motions of the artery added to the complexity of flow patterns in the PVS. Our study demonstrates that the combination of arterial wall expansion, rigid motions and a static CSF pressure gradient generates net and oscillatory PVS flow, quantitatively comparable with experimental findings. The static CSF pressure gradient required for net flow is small, suggesting that its origin is yet to be determined.

**Significance Statement:** Cerebrospinal fluid flow along perivascular spaces is hypothesized to be instrumental for clearance of metabolic waste from the brain, such as e.g. clearance of amyloid-beta, a protein known to accumulate as plaque within the brain in Alzheimer’s patients. Arterial pulsations have been proposed as the main driving mechanism for perivascular fluid flow, but it is unclear whether this mechanism alone is sufficient. Our results show that arterial pulsations drive oscillatory movement in perivascular spaces, but also indicate that a pressure gradient is required for net flow. However, the required pressure gradient is relatively small, thus suggesting that its origins can be associated with physiological processes within the brain and/or experimental procedures.

## Introduction

The glymphatic theory (1) suggests that the interaction of cerebrospinal fluid (CSF) and interstitial fluid facilitates the brain’s clearance of metabolites via perivascular spaces (PVS) in a process faster than diffusion alone. Experimental findings (2–7) demonstrate and support the concept of glymphatic function, while computational studies aiming to model the underlying mechanisms from first principles fail to show that perivascular coupling significantly accelerate transport when compared to diffusion (8–10). As a result, the glymphatic hypothesis is still controversial, almost a decade after its inception (11).

Perivascular flow appears to originate from forces associated with the cardiac cycle as travelling particles have a distinct cardiac frequency in their motion (2). The cardiac CSF pulsation is well characterized both in terms of CSF flow and intracranial pressures (ICP) (12). In humans, ICP is normally 7–15 mmHg (13, 14), and pulsates with a temporal peak-to-peak amplitude of around 1–4 mmHg (15, 16). The pulsation is almost synchronous within the whole cranium, yet there is a small spatial gradient of 1–5 mmHg/m (15, 17). Less data are available on the values of ICP and in particular pressure gradients in mice. Normal mouse ICP has been reported at 4 mmHg (18) with an approximate peak-to-peak temporal amplitude of 0.5–1 mmHg (19).

Forces inducing PVS flow may originate from local arterial expansions (2), but also from systemic ICP increase and blood pressure oscillations in proximal parts of the vasculature. The forces originating from systemic components are transmitted almost instantaneously to the PVS in terms of a pressure pulsation through the incompressible CSF. Peristalsis driven by the local arterial wall pulsation has received much attention, but computational modeling and theoretical calculations (in idealized geometries) point in different directions as to whether this mechanism is sufficient for net flow (10, 20–25).

In this study, we therefore address several forces with the potential to explain both net and oscillatory fluid movement in a realistic PVS geometry. In addition to the pulsatile local arterial expansion, we evaluated systemic CSF gradients of both static and pulsatile nature as well as rigid motions of the artery. We find that all forces combined may induce PVS flow comparable to experimental observations (2), but that the magnitude of the static pressure gradient required for net flow suggests that its origin is still unclear. A small net flow velocity close to the experimental data without the presence of a pressure gradient is only achieved when the PVS geometry is long (close to the wavelength of the arterial pulse).

## Results

To predict flow characteristics and detailed flow patterns in perivascular spaces surrounding pial arteries, we created a computational model of a CSF-filled PVS surrounding a bifurcating cerebral artery segment (Fig. 1A,B). Flow was induced in the PVS by combinations of local and systemic effects including pulsatile arterial wall motions, pulsatile arterial rigid motions, and static and/or pulsatile pressure differences between the inlet and outlets (Fig. 1C). We computed velocities and pressures in space and time, averaged normal velocities at the inlet and outlets over time, and net flow velocities (see Methods, Fig. 1D).

**Fig. 1.**
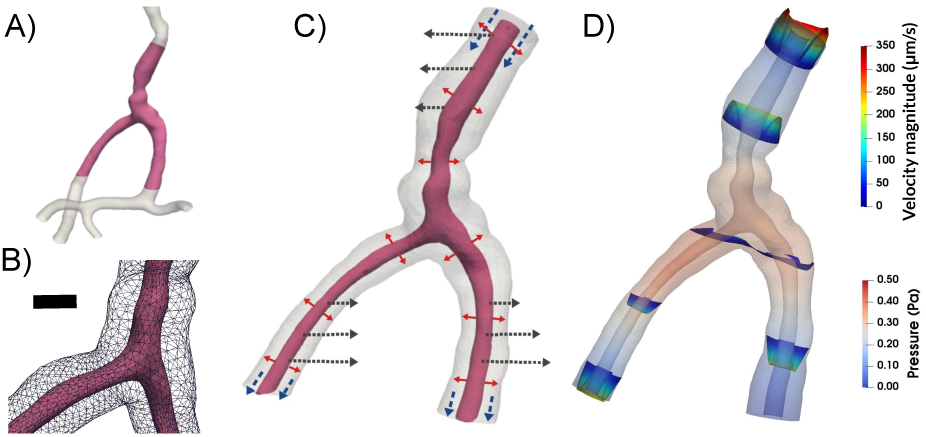
To study the mechanisms behind perivascular fluid flow, we extracted an image-based bifurcating arterial geometry **(A)** and generated a computational model of a surrounding perivascular space **(B)** subjected to different forces: arterial wall deformations (red arrows), systemic pressure variations (blue arrows) and rigid motions (black arrows) **(C)** to predict the induced CSF flow and pressure **(D)**. Scale bar: 0.05 mm.

### Vascular wall pulsations induce oscillatory bi-directional flow patterns in the PVS

When inducing flow in the PVS by pulsatile arterial wall motions, the fluid in the PVS oscillated with the same frequency (10 Hz) and in-phase with the wall. During systole flow was bi-directional: the arterial radius rapidly increased, pushing fluid out of the domain at both the inlet (top) and the outlets (bottom) (Fig. 2A). During diastole, flow was reversed (Supplementary Movie S1). Peak velocity magnitude occurred close to the inlet of the PVS model (Fig. 2A). From the inlet, the velocity magnitude decreased along the PVS until reaching a minimum close to the bifurcation. The velocity magnitude then increased towards the outlets but did not reach the same magnitude as at the inlet. The velocity profile in axial cross-sections of the PVS followed a Poiseuille-type flow pattern with high magnitude in central regions and low magnitude close to the walls (Fig. 2A). The (average normal) velocity at the inlet (see Methods) was negative (downwards into the PVS) during diastole reaching close to -45 *µ*m/s and positive (upwards, out of the PVS) during systole, reaching nearly 220 *µ*m/s, giving a peak-to-peak amplitude of 265 *µ*m/s (Fig. 2F).

**Fig. 2.**
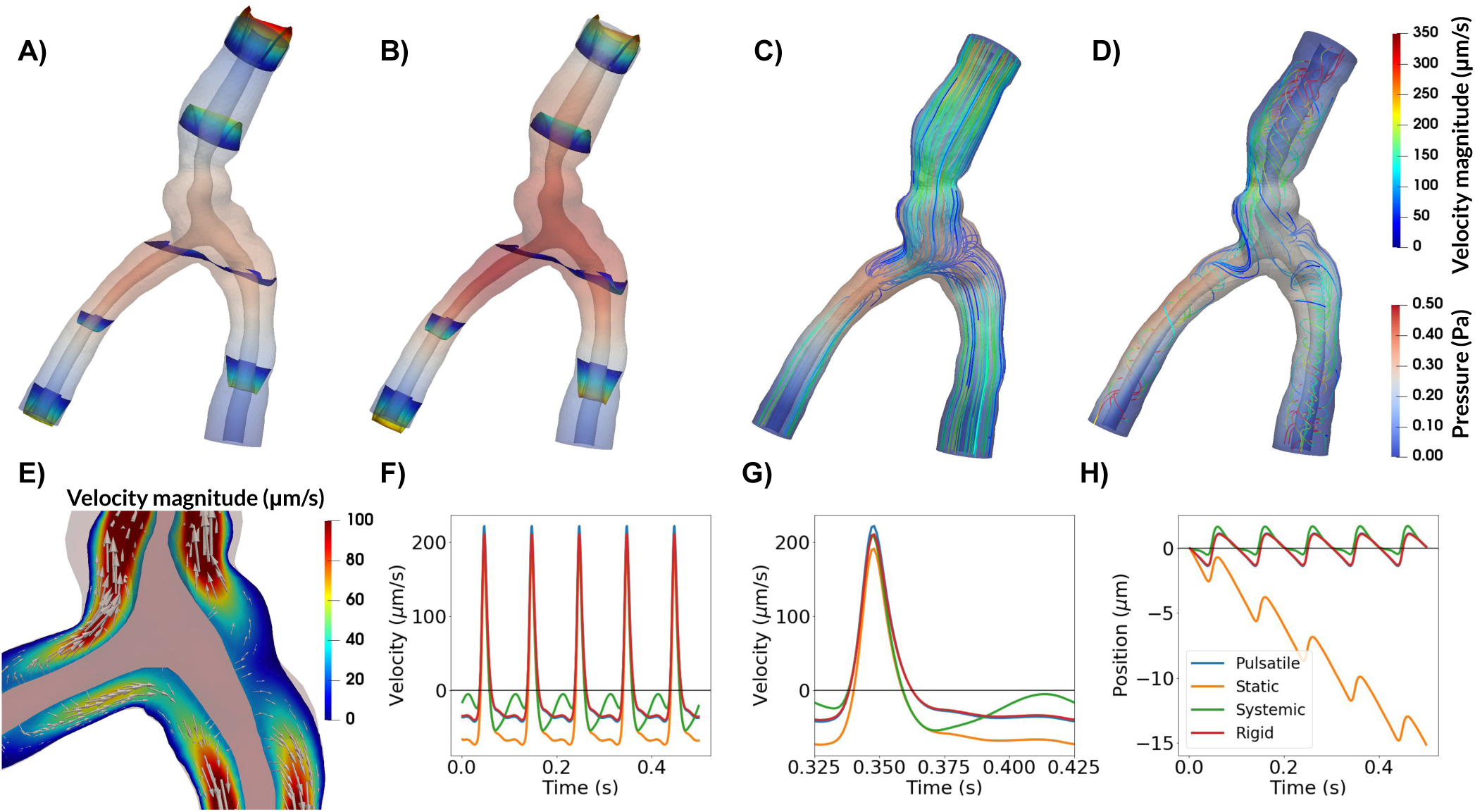
Top panel: Snapshots of CSF velocity and pressure at the time of peak velocity (or t = 0.048 s) for different types of forces. Pressure is shown everywhere with opacity, while velocity profiles are shown for given slices along the PVS (A, B) or as streamlines (C, D). **A)** With arterial wall deformations as the only driving force, pressure reaches a maximum close to the bifurcation, and velocity is stagnant at this point in space. As the artery expands, fluid flows in different directions within the PVS, always out of the domain, and velocities increase towards the inlet and outlet of the PVS. **B)** Adding a static pressure gradient of 1.46 mmHg/m (0.195 Pa/mm) results in higher peak velocities and less backflow, but oscillations due to arterial pulsations are still prominent. The pressure increases, but flow patterns are visually similar to (A). Differences can be seen in magnitude of flow at the inlets and outlets. **C)** A sinusoidally varying pressure gradient did not change the general flow pattern seen in (A, B) with parallel streamlines. **D)** Rigid motion of the artery caused less orderly CSF flow with more complex streamline patterns. **E)** Detailed flow patterns of the movement around bifurcations from (A). Slow flow close to the bifurcation is observed. **F)** Time profiles of the average normal velocity at the inlet for the four different models are similar, but differ in somewhat in shape during diastole. Negative values here correspond to flow downwards (in the direction of the net blood flow), while positive values correspond to flow upwards. **G)** Close-up on F) demonstrating that peak velocities are nearly identical for each model. **H)** Position plots of particles at the inlet indicating net flow only when a static pressure gradient is included. Net flow velocity at the inlet was 28*µ*m/s.

While the pressure difference between the PVS inlet and outlets was set at zero, the pressure within the PVS again oscillated with the cardiac frequency and varied almost linearly throughout the PVS (Fig. 2A). The peak pressure was 0.38 Pa (0.0029 mmHg), and occurred in the smallest daughter vessel after the bifurcation (≈ 0.36 mm from the left outlet). The time of peak pressure coincided with the time of peak velocity (t = 0.048 s). At time of peak pressure, the pressure gradient was nearly uniform throughout the domain, with an average gradient magnitude of 6.31 mmHg/m. High pressure gradients were observed locally in a small narrow region of the inlet vessel reaching a maximum gradient magnitude of 93.0 mmHg/m (data not shown).

### Static pressure gradients induce net PVS flow with backflow

Static CSF pressure gradients may occur as a direct consequence of tracer infusions (26). However, such gradients also occur naturally due to e.g. the third circulation (17, 27). To assess the static systemic effect and pulsatile local effect, we simulated flow and pressure under a static pressure difference between the inlet and outlets of the PVS model in combination with the pulsatile wall motion. Several pressure gradients were examined, representing forces involved in the cardiac and respiratory cycle, the third circulation, and in infusion tests (17, 26).

In general, the additional static pressure gradient induced net flow in the downwards direction, with the presence of oscillatory flow including backflow depending on the magnitude of the applied gradient and location. With a static pressure gradient of 1.46 mmHg/m (Fig. 2B), which is representative of the pulsatile pressure gradient induced by the cardiac cycle (17), net flow velocities varied between 20 and 30 *µ*m/s depending on the location of measurement: at the inlet, the net flow velocity was 28 *µ*m/s. The peak-to-peak amplitude of the average normal velocity was unchanged at 265 *µ*m/s (Fig. 2G). A particle suspended at the PVS inlet would then experience a pulsatile back-and-forth motion with a net movement downstream: 1-2 *µ*m upstream during systole and 5 *µ*m downstream during diastole (Fig. 2H). At the outlets, the average normal velocities were lower (in absolute value) due to the larger area and flow was nearly stagnant during diastole (backflow was negligible) (Supplementary Fig. S1). The static pressure gradient did otherwise not change the shape of the velocity pulse. We note that the pulsatile motion of particles was not easily visible, except for particles close to the inlet or outlets (Fig. 2)B).

The gradient induced by steady production of CSF i.e. corresponding to the third circulation (0.01 mmHg/m) generated negligible net flow, while a gradient equal to the respiratory gradient (0.52 mmHg/m) gave about threefold lower net flow velocity than the cardiac gradient described in detail above. An upper estimate of a gradient induced by infusion (≈12 mmHg/m) resulted in net flow velocity of more than 100 *µ*m/s.

### Flow induced by blood and CSF asynchrony

Flow in cranial or spinal PVS may be influenced by the relative timing of pulsatile blood and CSF pressures (28). With physiological pressure gradients, we investigated to what extent differences in phase – between the pulsatile arterial wall motion and some pulsatile systemic pressure gradient – would induce fluid velocities and net flow in the PVS (2, 3).

A pulsatile cardiac pressure gradient with peak amplitude 1.46 mmHg/m (=0.195 Pa/mm), frequency 10 Hz, combined with the pulsatile arterial wall motion at a phase shift *θ* (see Methods), again induced laminar flow in the PVS with streamlines taking the shortest path between the inlet and outlets (Fig. 2C). Further, the velocity profiles were comparable to those induced by the arterial wall motion alone (Fig. 2F–H, Supplementary Fig. S1). The velocity profile resulting from a phase shift of 10% of the cardiac cycle, while still similar, differed the most from the previous experiments. Qualitative differences could be observed during diastole, and the peak-to-peak amplitude of the velocity was slightly reduced at 262 *µ*m/s. However, no substantial net movement of fluid in any direction was observed: a particle suspended at the inlet of this systemic model would oscillate back-and-forth with a peak-to-peak amplitude of 2 *µ*m (Fig. 2H). Regardless of the phase shift applied in the systemic pressure gradient, net flow velocity did not exceed 0.5 *µ*m/s (Supplementary Fig. S2).

### Arterial rigid motions induce complex flow patterns

In addition to its pulsatile expansions and contractions, an artery can undergo pulsatile rigid motions i.e. rotations or local translations. The potential effect of such arterial rigid motions on PVS flow is poorly understood. To investigate, we extracted experimentally observed arterial rigid motions (2, Supplementary movie 2) and simulated how this additional movement could affect flow in the PVS (Fig. 3A).

**Fig. 3.**
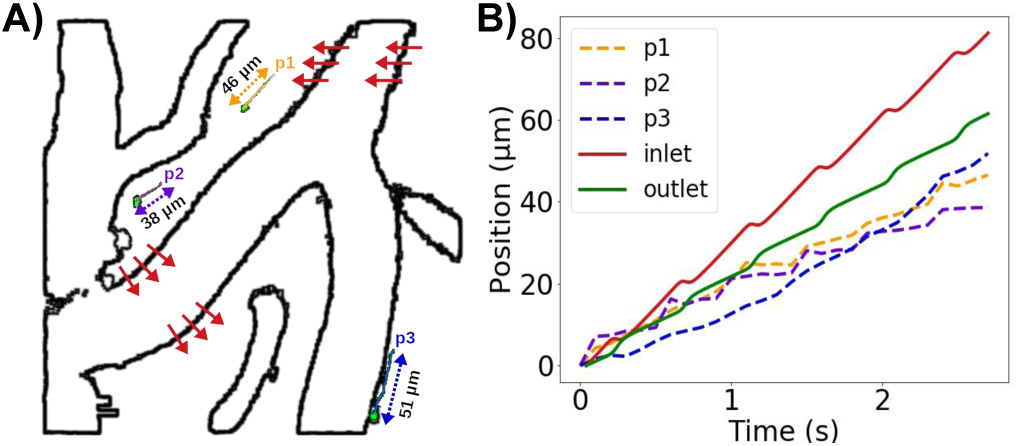
PVS flow predictions at a reduced arterial frequency and non-zero static pressure gradients compared with experimentally observed microsphere paths. **A)** Extraction of rigid arterial motions and three sample microspheres (p1, p2, p3) from experimental reports by Mestre et al (2). (Figure based on images adapted from (2, Supplementary movie 2) (CC BY 4.0)). Red arrows illustrate the rigid motion of the vessel with a peak amplitude of ≈ 6 *µ*m. **B)** Position (relative to each starting point) over time of model particles suspended at the PVS inlet and left outlet (2.2 Hz, static pressure gradient of 1.46 mmHg/m) compared to microsphere paths.

The rigid motions extracted were synchronous with the arterial wall pulsations. When combined with these pulsations, the rigid motion of the artery increased fluid motion within the PVS (Fig. 2D). As the artery shifted, the displaced fluid tended to move to the other side of the artery, thus yielding more complex streamline patterns and swirls. The peak-to-peak velocity amplitude was 251 *µ*m/s, which is slightly lower than for the other models. However, the rigid motion did not affect the overall shape of the velocity pulse at the inlet or outlets (Fig. 2F–G, Supplementary Fig. S1) or the net flow velocity (Fig. 2H). The rigid motion resulted in more complex flow, which will enhance local mixing.

### Arterial pulsation frequency modulates PVS flow velocity

The typical duration of the mouse cardiac cycle has been reported as 80-110 ms (29), corresponding to a cardiac frequency of 9–12.5 Hz. However, experimental studies of perivascular flow also also reveal cardiac frequencies as low as 2.2 Hz (2, Supplementary Movie 2). Reducing the frequency of the arterial wall pulsations from 10 to 2.2 Hz reduced the peak velocity by a similar factor: the peak-to-peak amplitude of the average normal velocity at the inlet was reduced from 260 to 60 *µ*m/s.

Adding a static gradient of 1.46 mmHg/m to the 2.2 Hz pulsations again induced net flow velocities of 20 - 30 *µ*m, but the pulsatile motion of particles was small compared to net flow (Supplementary Movie S2). Combining rigid motions with the experimentally observed arterial wall pulsation frequency of 2.2 Hz and a static pressure gradient of 1.46 mmHg/m, induced oscillatory PVS flow with non-trivial flow patterns, backflow, and net flow (Supplementary Movie S3, Fig. 3B). The pulsatile rigid motions induced oscillatory particle movement normal to the arterial wall, while the pulsatile arterial expansion induced back-and-forth movement along the PVS. Superimposed on the steady downwards flow induced by the static gradient, the movement of particles were thus similar to existing experimental observations (2) both in terms of net flow velocities and peak-to-peak pulsation amplitude.

### Model length modulates PVS velocity and net flow

Mathetal studies. On the other hand, a particle suspended at the matical modeling of PVS flow in idealized geometries has demonstrated that, under certain conditions, peristaltic motion of the arterial walls could induce substantial net flow velocities (22, 24). However, these findings have not been supported by computational models (30). The mathematical model (24) represents the PVS as an infinitely long annular cylinder, while in-vivo and in relevant computational models, the PVS is considerably shorter than the wavelength of the arterial pulse wave (30). To examine the effect of model length on PVS velocities and net flow, we considered a idealized axisymmetric model of an annular cylinder of different lengths *L* (1, 5, 10, 50 and 100 mm) for a fixed frequency (10 Hz) and culation (17) was not sufficient to drive net fluid movement arterial pulse wavelength *λ* (100 mm).

When the model length is shorter than the wavelength, velocities are highly dependent on the length of the PVS (Fig. 4). For *L* ≪ *λ*, the wall displacement is close to uniform along the PVS, and more fluid will leave the domain through the inlet and outlet in a longer artery (Fig. 4C). Thus, for a given relative wall displacement and model lengths smaller than half the arterial pulse wavelength, the velocity at the inlet (or outlet) will increase with increasing PVS model length (Fig. 4A). The shape of the velocity pulse also change: for longer models, at the inlet, we observe a longer period of upwards flow (out of the domain) and a corresponding shorter period of downwards flow (into the domain). When the domain length is equal to an integer multiple of the wavelength, using a zero pressure drop or a symmetry boundary condition will model an infinitely long cylinder. For this case, the velocity will not increase further with increasing PVS length (data not shown). To obtain a peak-to-peak velocity amplitude of ≈20 *µ*m/s (2), a frequency of 10 Hz required a PVS length of 0.10 mm. Changing the frequency to 2.2 Hz, required a PVS length of 0.47 mm to reach the same amplitude (data not shown).

**Fig. 4.**
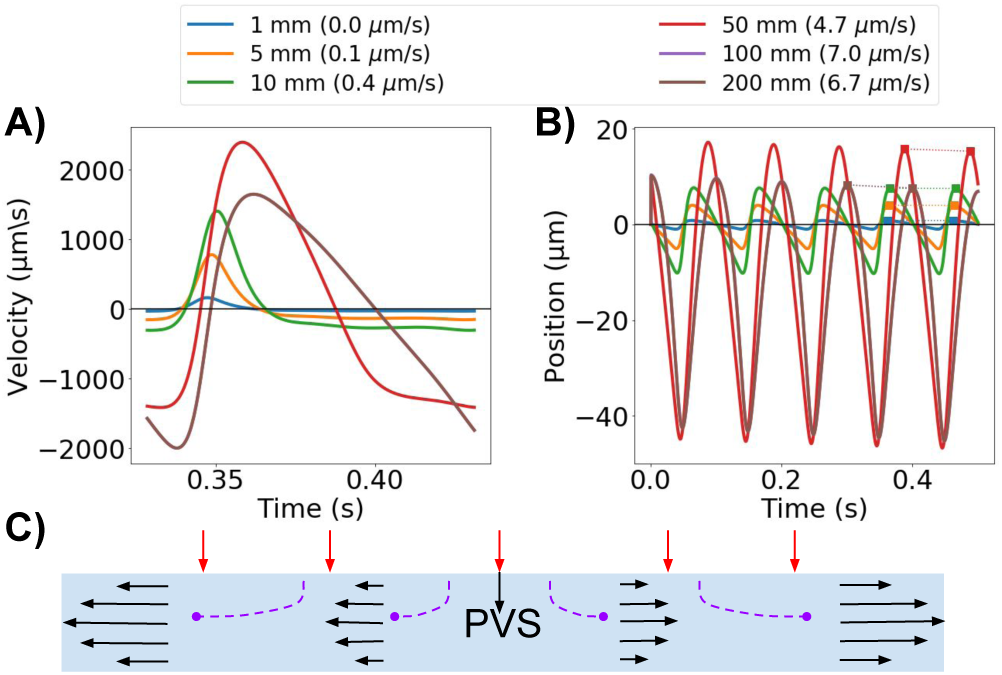
PVS model length modulates velocity. **A)** The average normal velocity at the inlet in idealized PVS models increases with increasing model length. **B)** Position of a particle moving with the velocity at the inlet. Net flow velocities (in parenthesis for each model length in the legend) are small compared to the large average normal velocity amplitudes, but can reach up to 7 *µ*m/s for the longer models (100 mm). **C)** Schematic of the idealized axisymmetric model. For long wavelengths, the displacement is almost uniform along the PVS. Increased PVS length will thus increase velocity at the model ends as more fluid needs to escape the domain.

For the same set of geometries, net flow also increased with model lengths up to the wavelength, and in the long idealized models of lengths 50 and 100 mm, net flow velocities of 4.7 and 7.0 *µ*m/s were observed (Fig. 4B). For the other PVS lengths tested, the net flow velocity was lower than 1 *µ*m/s. Net flow velocities were small compared to the large average normal velocity amplitudes: a particle suspended at the PVS inlet could experience a change in position of up to 60 *µ*m over one cardiac cycle (10 Hz, model length 50 mm) (Fig. 4B).

## Discussion

Experimental studies of perivascular flow have found substantial velocities and net particle movement, predominantly in a uni-directional pattern. Mestre et al (2) reported a peak-to-peak velocity amplitude of ≈20 *µ*m/s and a typical net flow velocity of 18.7 *µ*m/s. With a period of 0.45 s (2.2 Hz), a particle could then be expected to move no more than 9 *µ*m back-and-forth per cycle. Similarly, Bedussi et al (3) reported an average net flow velocity of 17 *µ*m/s and a mean amplitude of movement of 14 *µ*m. However, the shorter cardiac period of 0.15 s points at much higher velocity amplitudes (at least 100-200 *µ*m/s) in the latter study. Our peak-to-peak velocity amplitudes (251-265 *µ*m/s for 10 Hz, 60 *µ*m/s for 2.2 Hz) are thus at the upper range of experimental values reported. Our observations further point at the impact of cardiac frequency on velocity amplitude, which may explain the difference in estimated velocity amplitudes between these two experimental studies. On the other hand, a particle suspended at the PVS inlet in our model would oscillate back-and-forth with a peak-to-peak amplitude of 2 *µ*m, with an additional net downstream movement only if a static pressure gradient is imposed. These changes in position are thus at the lower end of the experimental observations, pointing at the likely presence of a static CSF pressure gradient in the experimental configurations.

A static CSF pressure gradient – of magnitude corresponding to the pulsatile gradient induced by the cardiac cycle – was sufficient to create net flow velocities of 30–40 *µ*m/s in the PVS. A pressure gradient associated with the third circulation (17) was not sufficient to drive net fluid movement in the PVS. The respiratory gradient is approximately one third of the cardiac gradient (17), and was sufficient to drive some PVS flow when applied as a static pressure difference. Thus, longer waves (such as those induced by respiration and vasomotion (31)) may also play a role in net fluid movement in the PVS. Our upper estimate of the pressure gradient induced between the CSF and the PVS during an infusion test (in humans) (26) resulted in net flow velocities of several hundred *µ*m/s, much higher than what has been observed in mice (2, 3). Overall, our observations indicate that the static pressure gradient sufficient to drive net flow is small (≈1.5 mmHg/m, i.e. a pressure difference of 0.015 mmHg per cm) compared to the intracranial pressure increase of 1.4–3 mmHg observed in mice during tracer infusion (18, 32).

Arterial rigid motions were of greater amplitude than the arterial wall expansions, but had minimal effect on average and net flow velocities. Indeed, the rigid motions of the artery did not force fluid to leave the PVS domain, but rather displaced fluid within. As such, arterial rigid motions can create oscillatory movement of cardiac frequency within the PVS without adding to the net movement of particles. In our model, rigid motions and arterial expansion combined can explain oscillatory motion as seen by Mestre et al. (2), but were not sufficient to generate net flow. However, the rigid motion introduced complex swirling that would significantly enhance local mixing and potentially contribute to increased clearance.

A systemic CSF pulsation of physiological amplitude (17, 26) – out of synchrony with the arterial wall pulsation – did not induce net fluid movement in the brain PVS. The relative timing of arterial and CSF pulse waves, a possible factor for net fluid movement in spinal cord PVS (28), is thus not a likely factor for explaining higher average or net flow velocities in brain PVS.

The length of the PVS segment is important for the observed fluid dynamics in the domain, and is also an important modeling parameter. When the PVS model is much shorter than the wavelength of the arterial pulse wave, the velocity amplitudes at the inlet and outlets of the PVS are directly linked to the PVS length. Similar effects have previously been noted by Asgari et al. (10) as an increase in dispersion effects with length, and by Rey and Sarntinoranont (25) as a variable Péclet number throughout the PVS domain. Considerable net flow in our models were seen only for very long geometries.

Several modelling studies have now tried to explain net movement of fluid in the PVS driven by local arterial pulsations. While theoretical considerations have explained net flow by arterial wall pulsations alone (22, 24), most computational studies (10, 20, 25, 30) suggest that the local effect of arterial wall pulsations is not sufficient to drive net flow in the PVS of magnitude comparable to experimental observations. Theoretical considerations have assumed an infinitely long cylinder, which we here show overestimates net flow and velocity amplitudes compared to PVS models of physiologically relevant lengths. In our idealized computational models however, net flow velocity is higher than the theoretical model by Wang and Olbricht (24) predicts: with a 0.7% half-amplitude of the arterial wall expansion, inner and outer radii of 20 and 60 *µ*m, and an arterial wave speed of c = 1 m/s, their model predicts an average net flow velocity of 1.53 *µ*m/s, which is lower than our estimates of 6.7–7 *µ*m/s by a factor of four. It should be emphasized that while the Wang and Olbricht net flow model (24) does not differentiate between PVS lengths, it is sensitive to PVS width and half-amplitude; thus specific such parameters could yield a higher net flow velocity.

In terms of limitations, PVS flow was modelled as incompressible viscous fluid flow ignoring potential barriers to flow (reduced permeability) in contrast to e.g. (23). For pial PVS, this may be a reasonable assumption. The addition of a finite permeability would be expected to yield lower velocities. We also ignored nonlinear (turbulent) effects, which, given the low Reynolds numbers involved, seems a reasonable approximation. The PVS domain was assumed to have a constant cross-sectional width, while other studies have suggested that the PVS cross-section is elliptic (2, 33). For the representation of the rigid motion, we estimated its magnitude (≈6 *µ*m) without isolating the ≈1 *µ*m wall pulsations. The findings reported here thus represents an upper estimate of the impact of arterial rigid motions. Finally, all pressure gradients used to drive flow in our models originated from human measurements, while both the PVS size and previously reported velocities were obtained in mice. Compared to humans, mice have smaller CSF volumes, lower ICP and ICP amplitudes and a shorter CSF turnover time (34). The latter suggests that the third circulation gradient is larger in mice than in humans. However, an increase of a factor ≈50 from human to mouse would be needed for the third circulation gradient to drive any substantial net flow. We mainly considered static pressure gradients, but other non-zero cardiac cycle averaged pressure gradients could yield similar results.

In conclusion, our simulations indicate that the combination of arterial wall pulsations and a systemic static pressure gradient larger than that associated with the third circulation can explain experimental findings on pulsatile perivascular flow. The required static gradient need not necessarily be caused mainly by an infusion, as such a gradient is at the order of physiological pressure gradients in the brain. Without a pressure gradient, net flow was only achieved for very long PVS geometries (on the order of the wavelength, here 100 mm), explaining why theoretical considerations of infinitely long cylinders yield net flow. Finally, rigid arterial motion can induce complex flow patterns in the PVS.

## Materials and Methods

### PVS geometry and mesh generation

The PVS geometry was generated from model representations of cerebral arteries (case id C0075) from the Aneurisk dataset repository (35). The full geometry was clipped to define a vessel segment including the arterial bifurcation with one inlet vessel and two outlet vessels (Fig. 1A, Supplementary Fig. S3). The PVS domain was defined by creating an annular cylinder surrounding the artery with the arterial wall as its inner surface. The width of each PVS was set proportional to the arterial diameter and scaled to the mouse scale. The PVS center line was then of maximal length 1 mm with inlet and outlet branches of comparable lengths (≈0.5 mm), PVS widths of 28–42 *µ*m and inner arterial diameters of 32–46 *µ*m. We created a finite element mesh of the PVS with 174924 tetrahedrons and 34265 vertices using VMTK (36).

We also defined a set of idealized PVS domains as annular cylinders of lengths *L* ∈ [1, 5, 10, 50, 100] mm with a annular crosssection width of 40*µm*. These annular cylinders were represented by one-dimensional axisymmetric finite element meshes with 10*L* + 1 vertices (mesh size 0.1 mm).

### CSF flow model and parameters

To model the flow of CSF in the pial PVS, we consider the CSF as an incompressible, viscous fluid flowing at low Reynolds numbers in a moving domain – represented by the time-dependent Stokes equations over a time-dependent domain. The initial PVS mesh defines the reference domain Ω_0_ for the CSF with spatial coordinates *X ∈* Ω_0_. We assume that the PVS domain Ω_*t*_ at time *t >* 0 has spatial coordinates *x ∈* Ω_*t*_ and is defined as a deformation of the reference domain Ω_0_ ↦Ω_*t*_ with *x* = *d*(*X, t*) for a prescribed space- and time-dependent domain deformation *d* with associated domain velocity *w*.

The fluid velocity *v* = *v*(*x, t*) for *x ∈* Ω_*t*_ at time *t* and the fluid pressure *p* = *p*(*x, t*) then solves the following system of time-dependent partial differential equations (PDEs) (37):

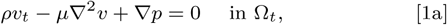

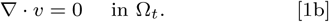

The CSF dynamic viscosity *µ* is set to 0.697 Pa/s with density *ρ* = 10^3^ kg/m^3^. On the inner PVS wall we set the CSF velocity *v* to match the given domain velocity *w*. We assume that the outer PVS wall is impermeable and rigid with zero CSF velocity. At the PVS inlet and outlet, we impose given pressures in the form of traction conditions. The system starts at rest and we solve for a number of flow cycles to reach a periodic steady state.

### Pulsatile wall motion and velocity

We stipulate that arterial blood flow pulsations induce a pulsatile movement of the inner PVS boundary Λ. We let this boundary deform in the direction of the boundary normal with a spatially and temporally varying amplitude *A*:

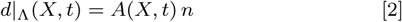

where *n* denotes the outward pointing boundary normal. For the amplitude *A*, we consider the combination of (i) the wall motions reported by Mestre et al (2, Fig. 3e–f) and (ii) a travelling wave along the PVS length. We extracted the percentage change in artery diameter Δ*d/d*(*s*) as a function of the fraction of the cardiac cycle *s* (2, Fig. 3e) using WebPlotDigitizer (38). To represent the spatial variation, we assume that the arterial pulse wave takes the form of a periodic travelling wave with wave speed *c* = 1 m/s (2) (and corresponding wave length *λ* = *c/f* for a given cardiac frequency *f*). We then set

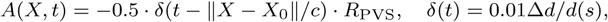

where *s* is the fraction of the cardiac cycle: *s* = (*t · f*) mod 1, with an average PVS width *R*_PVS_ = 4.4 *×* 10^*−*2^ mm, and *X*_0_ is a fixed point close to the center of the inlet. We let the default cardiac period be 1*/f* = 0.1 s (frequency 10 Hz).

### Static and pulsatile pressure gradients

Pressure gradients in the arterial PVS can also be a consequence of the systemic phase-shift in pressure pulsations between larger proximal arteries, CSF, and the venous system (39), a general pressure increase in the CSF due to infusion (26), or other factors affecting the relative timing of the arterial and CSF pulse pressure (28). To examine these systemic effects, we considered different static and pulsatile pressure gradients.

We associated static pressure gradients with the third circulation (27) (0.01 mmHg/m), the cardiac cycle (1.46 mmHg/m), and respiration (0.52 mmHg/m). The latter two values correspond to the peak amplitude of the pulsatile pressure gradients associated with these cycles (17, 40), and should be considered as upper estimates of any associated static pressure gradients. During an infusion a change of at least 0.03 mmHg may occur between the CSF and the PVS (26). The pressure drop occurs over at least a cortex thickness of 2.5 mm (41), and an upper estimate of the static pressure gradient due to infusion can thus be computed as

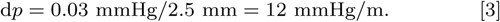

Inspired by Bilston et al (28), we also considered a cardiac-induced pulsatile pressure difference between the inlet and outlet with a phase shift relative to the pulsatile arterial wall movement of the form

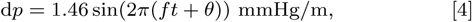

where *f* is the pulsatile frequency, and *θ* is the relative phase shift ranging from 0 to 1.

The pressure gradients where weighted by the lengths of each branch to ensure that average pressure gradients from the inlet to the two different outlets are equal, and then applied as pressure differences between the inlet and outlets as traction boundary conditions.

### Image analysis of rigid motion and particle positions

We defined the arterial rigid motions by juxtaposing 28 screenshots, extracted at a fixed frequency, from (2, Supplementary movie 2). Comparing the artery outlines and motion, we estimated the peak amplitude *γ* of the motion to be no more than 6 *µ*m (Fig. 3A) and identified a center point for the rigid motion *X*_*c*_ close to the center of the bifurcation. We then defined the signed amplitude of the rigid motion as

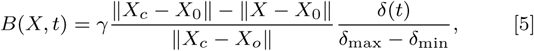

where *δ*_min_ (resp. *δ*_max_) is the minimum (resp. maximum) relative diameter variation, and ‖*X*_*c*_ − *X*_0_‖ ≈ 0.40 mm. We defined the (normalized) direction of the rigid motion *r* as normal to the main axis between *X*_0_ and *X*_*c*_. We then investigated the impact of rigid motion on perivascular flow by imposing the following boundary displacement (in place of Eq. (2)):

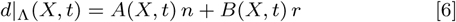

From the screenshots, we also tracked the position of a number of sample microspheres over time using GIMP-Python (42).

### Numerical solution

To compute numerical solutions of Eq. (1), we consider the Arbitrary Lagrange-Eulerian (ALE) formulation (37) with a first order implicit Euler scheme in time and a second-order finite element scheme in space. For each discrete time *t*^*k*^ (*k* = 1, 2, *…*), we evaluate the boundary deformation *d*|_Λ_ given by Eq. (2) and extend the deformation to the entire mesh by solving an auxiliary elliptic PDE. The computational mesh is deformed accordingly and thus represents 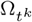. We also evaluate the first order piecewise linear discrete mesh velocity 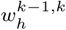 associated with this mesh deformation.

Next, at each discrete time *t*^*k*^, we solve for the approximate CSF velocity 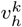 and pressure 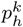 on the domain 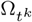 satisfying

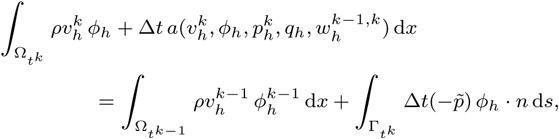

where

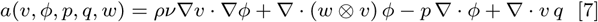

for finite element test functions *ϕ*_*h*_ and *q*_*h*_ defined on 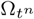, Δ*t* is the time step size, 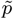 is the prescribed boundary pressure at the inlet and/or outlet 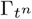, and 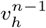 and 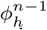 are the approximate velocity and test function respectively at the previous discrete time *t*^*n−*1^, defined over the previous domain Ω_*t*_*n−*1. We set Δ*t* = 0.001 s.

### Computation of output quantities

With the computed velocity field *v*, we define the flow rate at the inlet 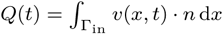, and the average normal velocity (see e.g. Fig. 2G) was computed as *v*_avg_(*t*) = *Q*(*t*)*/A*_in_, where *A*_in_ is the area of the inlet. From the average normal velocity, the position of a particle at time *t* was computed as

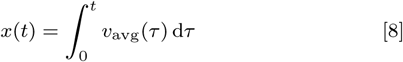

Finally, the net flow velocity was computed as the slope between the peaks of *x*(*t*), using the two last cardiac cycles.

### Computational verification

All numerical results were computed using the FEniCS finite element software suite (43). Key output quantities were compared for a series of mesh resolutions and time step sizes to confirm the convergence of the computed solutions (Supplementary Fig. S4). The simulation code, meshes and associated data are openly available (44).

## ACKNOWLEDGMENTS

This research is supported by the European Research Council (ERC) under the European Union’s Horizon 2020 research and innovation programme under grant agreement 714892, and by the NOTUR grant NN9316K. The third author was supported by the Research Council of Norway under grant agreement 301013.

The authors declare no competing interests.

## Supplementary information

**Fig. S1.**
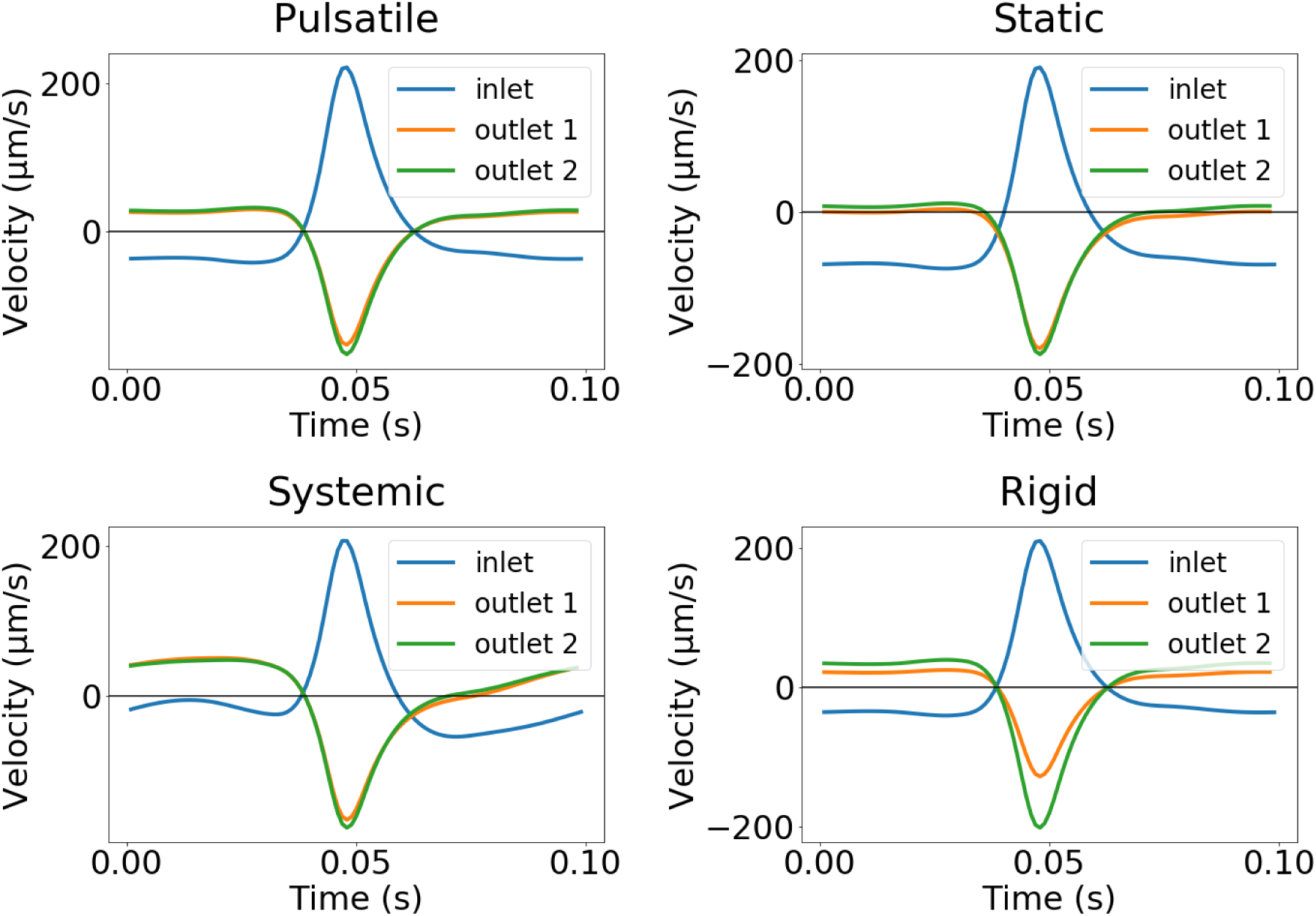
Comparison of average normal velocity at the inlet and outlet for each model: pulsatile arterial wall motion only (Pulsatile), with an additional static pressure gradient associated with the cardiac cycle (Static), with an additional pulsatile pressure gradient (Systemic), or with an additional rigid motion (Rigid). Cardiac frequency: 10 Hz. The outward pointing normal at the inlet defines the positive direction; negative values thus refer to flow downwards (into the PVS at the inlet, and out of the PVS at the outlet). All models predict bi-directional flow during systole, with fluid leaving the domain at both inlet and outlets. The peak velocity amplitude is slightly higher at the inlet during systole mainly due to the smaller area for flow compared to the combined area at the outlets. The velocity during diastole differ more between the models, and more (in absolute value) between at the inlet and outlet.

**Fig. S2.**
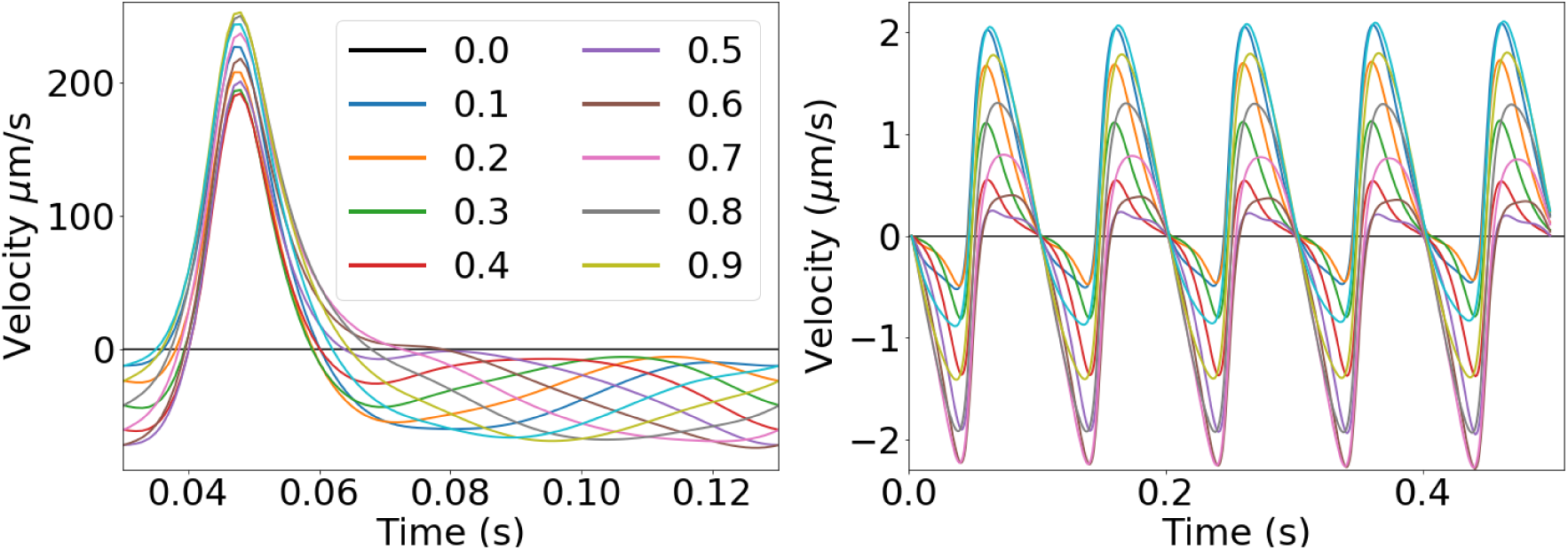
Comparison of average normal velocities for the model with a pulsatile pressure gradient with different phase shifts between the systemic CSF pressure and the arterial wall pulse wave. The shift *θ* was represented by different fractions of the cardiac cycle ranging from 0 to 0.9 of steps 0.1. The net flow velocity was lower than 0.5 *µ*m/s for all shifts *θ*.

**Fig. S3.**
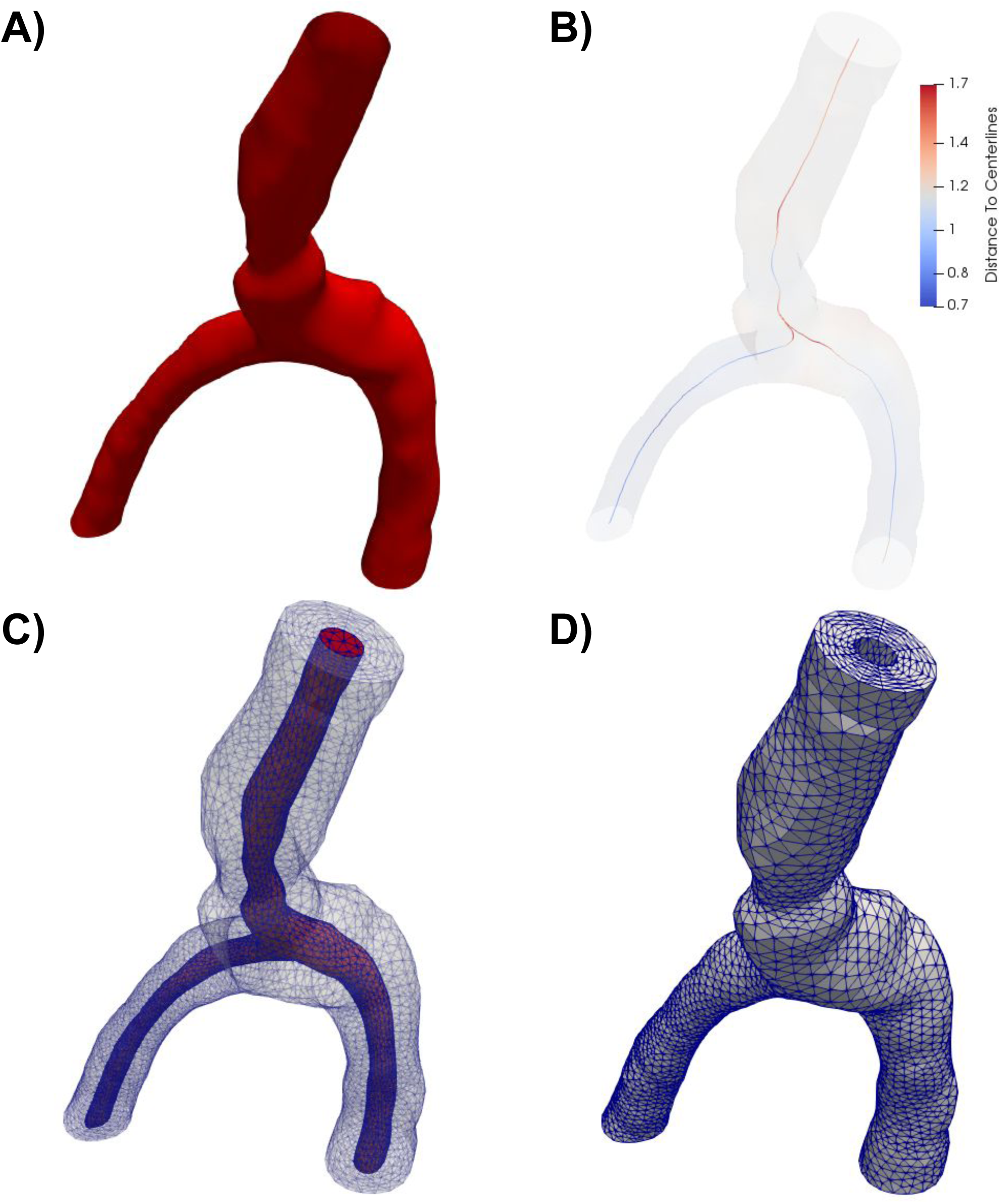
Overview of the computational mesh generation. **A)** The artery geometry was extracted from Aneurisk dataset repository (case id C0075) (35). **B)** The domain center line used to determine the PVS width was computed using VMTK. The color indicates the distance from the center line to the vessel wall. A finite element mesh was generated of the full geometry (including both the artery and the PVS) (**C**), before the PVS mesh (**D**) was extracted for simulations.

**Fig. S4.**
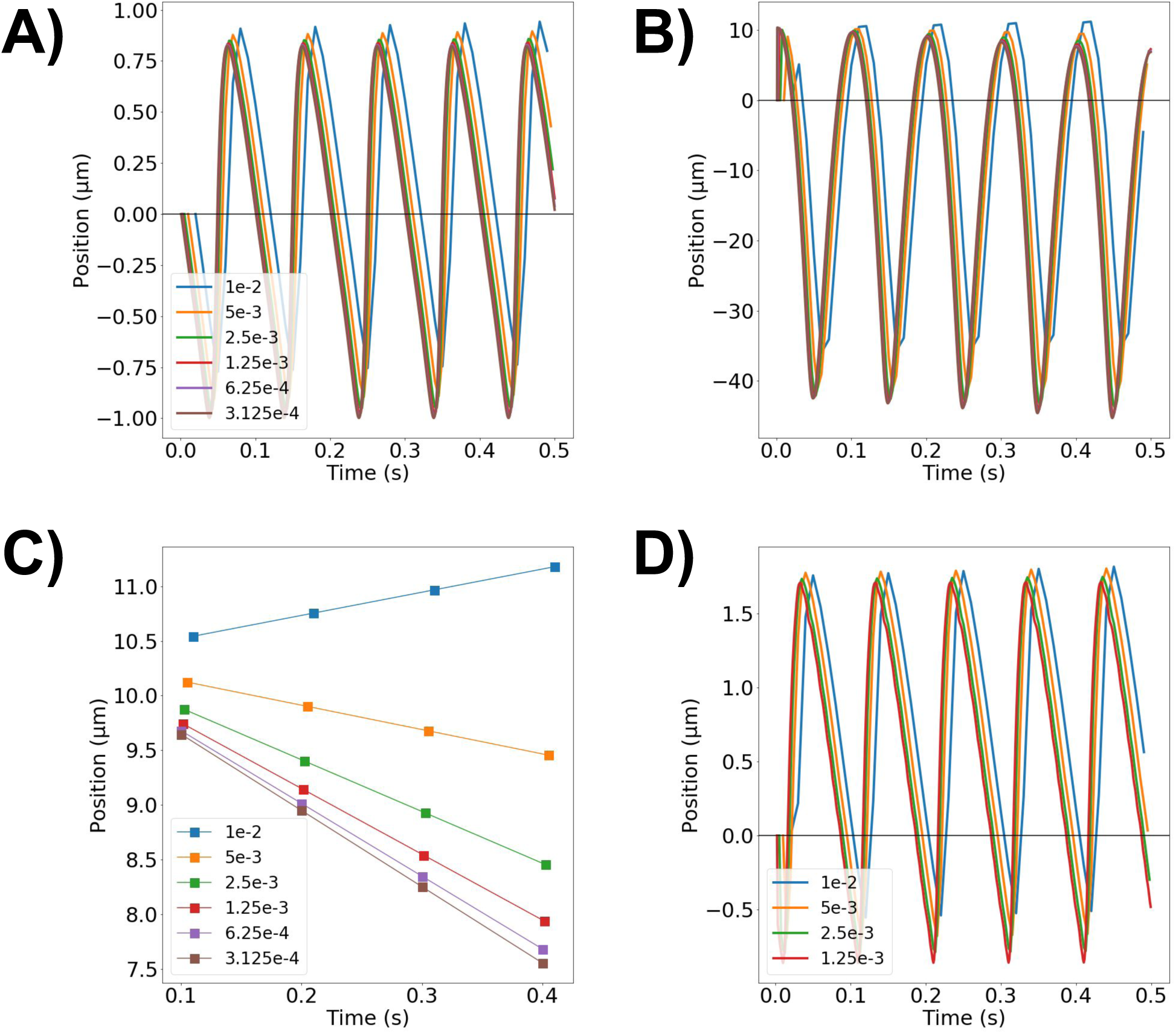
Numerical verification and convergence analysis of the computational models. Position over time for the idealized models of length 1 mm (**A**) and 100 mm (**B**) for different time resolutions (Δ*t*). Key output quantities converge as the time resolution is reduced as expected. For a PVS length of 100 mm, the time step required for convergence was lower than for the shorter models. **C**) Peaks of the position over time for different time resolutions (L = 100 mm). We note that the computed net flow velocity strongly depends on the time resolution, but that a time step of 1 ms is sufficient. **D**) Position over time for the bifurcating arterial geometry (C0075) for different time resolutions. The time step of 1 ms is again sufficient.

## References

1. JJ Iliff, et al., A paravascular pathway facilitates CSF flow through the brain parenchyma and the clearance of interstitial solutes, including amyloid-*β*. Sci. translational medicine 4, 147ra111–147ra111 (2012).

2. H Mestre, et al., Flow of cerebrospinal fluid is driven by arterial pulsations and is reduced in hypertension. Nat. communications 9, 4878 (2018).

3. B Bedussi, M Almasian, J de Vos, E VanBavel, EN Bakker, Paravascular spaces at the brain surface: Low resistance pathways for cerebrospinal fluid flow. J. Cereb. Blood Flow & Metab. 38, 719–726 (2018).

4. MJ Hannocks, et al., Molecular characterization of perivascular drainage pathways in the murine brain. J. Cereb. Blood Flow & Metab. 38, 669–686 (2018).

5. Q Ma, et al., Rapid lymphatic efflux limits cerebrospinal fluid flow to the brain. Acta neuropathologica 137, 151–165 (2019).

6. G Ringstad, et al., Brain-wide glymphatic enhancement and clearance in humans assessed with MRI. JCI insight 3 (2018).

7. L Xie, et al., Sleep drives metabolite clearance from the adult brain. Science 342, 373–377 (2013).

8. KE Holter, et al., Interstitial solute transport in 3D reconstructed neuropil occurs by diffusion rather than bulk flow. Proc. Natl. Acad. Sci. 114, 9894–9899 (2017).

9. JH Smith, JA Humphrey, Interstitial transport and transvascular fluid exchange during infusion into brain and tumor tissue. Microvasc. research 73, 58–73 (2007).

10. M Asgari, D De Zélicourt, V Kurtcuoglu, Glymphatic solute transport does not require bulk flow. Sci. reports 6, 38635 (2016).

11. NJ Abbott, ME Pizzo, JE Preston, D Janigro, RG Thorne, The role of brain barriers in fluid movement in the CNS: is there a ‘glymphatic’ system? Acta neuropathologica 135, 387–407 (2018).

12. ME Wagshul, PK Eide, JR Madsen, The pulsating brain: a review of experimental and clinical studies of intracranial pulsatility. Fluids Barriers CNS 8, 5 (2011).

13. J Malm, J Jacobsson, R Birgander, A Eklund, Reference values for CSF outflow resistance and intracranial pressure in healthy elderly. Neurology 76, 903–909 (2011).

14. M Czosnyka, JD Pickard, Monitoring and interpretation of intracranial pressure. J. Neurol. Neurosurg. & Psychiatry 75, 813–821 (2004).

15. G Ringstad, et al., Non-invasive assessment of pulsatile intracranial pressure with phase-contrast magnetic resonance imaging. PloS one 12, e0188896 (2017).

16. PK Eide, E Kerty, Static and pulsatile intracranial pressure in idiopathic intracranial hypertension. Clin. neurology neurosurgery 113, 123–128 (2011).

17. V Vinje, et al., Respiratory influence on cerebrospinal fluid flow–a computational study based on long-term intracranial pressure measurements. Sci. reports 9, 9732 (2019).

18. H Mestre, Y Mori, M Nedergaard, The brain’s glymphatic system: Current controversies. Trends Neurosci. xx, 1–9 (2020).

19. M Uldall, et al., A novel method for long-term monitoring of intracranial pressure in rats. J. neuroscience methods 227, 1–9 (2014).

20. AD Martinac, L. Bilston, Computational modelling of fluid and solute transport in the brain. Biomech. modeling mechanobiology xx, 1–20 (2019).

21. AK Diem, et al., Arterial pulsations cannot drive intramural periarterial drainage: significance for A*β* drainage. Front. neuroscience 11, 475 (2017).

22. JH Thomas, Fluid dynamics of cerebrospinal fluid flow in perivascular spaces. J. Royal Soc. Interface 16, 20190572 (2019).

23. MK Sharp, RO Carare, BA Martin, Dispersion in porous media in oscillatory flow between flat plates: applications to intrathecal, periarterial and paraarterial solute transport in the central nervous system. Fluids Barriers CNS 16, 13 (2019).

24. P Wang, WL Olbricht, Fluid mechanics in the perivascular space. J. theoretical biology 274, 52–57 (2011).

25. J Rey, M Sarntinoranont, Pulsatile flow drivers in brain parenchyma and perivascular spaces: a resistance network model study. Fluids Barriers CNS 15, 20 (2018).

26. V Vinje, A Eklund, KA Mardal, ME Rognes, K. Støverud, Intracranial pressure elevation alters CSF clearance pathways. Fluids Barriers CNS 17, 1–19 (2020).

27. H Cushing,, et al., The third circulation and its channels. Lancet 2, 851–857 (1925).

28. LE Bilston, MA Stoodley, DF Fletcher, The influence of the relative timing of arterial and subarachnoid space pulse waves on spinal perivascular cerebrospinal fluid flow as a possible factor in syrinx development. J. neurosurgery 112, 808–813 (2010).

29. S Kaese, S Verheule, Cardiac electrophysiology in mice: a matter of size. Front. Physiology 3, 345 (2012).

30. R Kedarasetti, PJ Drew, F Costanzo, Arterial pulsations drive oscillatory flow of CSF but not directional pumping. bioRxiv xx, xx (2020).

31. RO Carare, et al., Vasomotion drives periarterial drainage of a*β* from the brain. Neuron 105, 400–401 (2020).

32. JJ Iliff, et al., Cerebral arterial pulsation drives paravascular CSF–interstitial fluid exchange in the murine brain. J. Neurosci. 33, 18190–18199 (2013).

33. J Tithof, DH Kelley, H Mestre, M Nedergaard, JH Thomas, Hydraulic resistance of periarterial spaces in the brain. Fluids Barriers CNS 16, 19 (2019).

34. WM Pardridge, CSF, blood-brain barrier, and brain drug delivery. Expert. opinion on drug delivery 13, 963–975 (2016).

35. Aneurisk-Team, AneuriskWeb project website, http://ecm2.mathcs.emory.edu/aneuriskweb (Web Site) (2012).

36. L Antiga, et al., An image-based modeling framework for patient-specific computational hemodynamics. Med. & Biol. Eng. & Comput. 46, 1097–1112 (2008).

37. J San Martín, L Smaranda, T Takahashi, Convergence of a finite element/ALE method for the Stokes equations in a domain depending on time. J. computational applied mathematics 230, 521–545 (2009).

38. A Rohatgi, WebPlotDigitizer (2017).

39. P Holmlund, et al., Venous collapse regulates intracranial pressure in upright body positions. Am. J. Physiol. Integr. Comp. Physiol. 314, R377–R385 (2018).

40. PK Eide, T Sæhle, Is ventriculomegaly in idiopathic normal pressure hydrocephalus associated with a transmantle gradient in pulsatile intracranial pressure? Acta neurochirurgica 152, 989–995 (2010).

41. B Fischl, AM Dale, Measuring the thickness of the human cerebral cortex from magnetic resonance images. Proc. Natl. Acad. Sci. 97, 11050–11055 (2000).

42. GNU Image Manipulation Program, version 2.8.22 (2017).

43. C. M Alnæs, et al., The FEniCS project version 1.5. Arch. Numer. Softw. 3 (2015).

44. C Daversin-Catty, V Vinje, KA Mardal, ME Rognes, mechanisms-behind-pvs-flow-v1.0 (source code) (2020).

